# Dual Passive Reactive Brain Computer Interface: a Novel Approach to Human-Machine Symbiosis

**DOI:** 10.1101/2021.12.23.473161

**Authors:** Frédéric Dehais, Simon Ladouce, Ludovic Darmet, Nong Tran-Vu, Giuseppe Ferraro, Juan Torre Tresols, Sébastien Velut, Patrice Labedan

## Abstract

The present study proposes a novel concept of neuroadaptive technology, namely a dual passive-reactive Brain-Computer Interface (BCI), that enables bi-directional interaction between humans and machines. We have implemented such a system in a realistic flight simulator using the NextMind classification algorithms and framework to decode pilots’ intention (reactive BCI) and to infer their level of attention (passive BCI). Twelve pilots used the reactive BCI to perform checklists along with an anti-collision radar monitoring task that was supervised by the passive BCI. The latter simulated an automatic avoidance maneuver when it detected that pilots missed an incoming collision. The reactive BCI reached 100% classification accuracy with a mean reaction time of 1.6*s* when exclusively performing the checklist task. Accuracy was up to 98.5% with a mean reaction time of 2.5*s* when pilots also had to fly the aircraft and monitor the anti-collision radar. The passive BCI achieved a *F*_1_ − score of 0.94. This first demonstration shows the potential of a dual BCI to improve human-machine teaming which could be applied to a variety of applications.

## 1 INTRODUCTION

Brain-Computer Interfaces (BCI) offer a direct communication pathway between a user and a machine without requiring any muscular engagement (Clerc et al., 2016). To this end, BCI derives the user’s intentions and mental states from neural signals. The decoding of specific neural signals triggers interactions with aspects of the computerized environment (*e.g*., moving a cursor, keystrokes) or external devices (*e.g*. prostheses) to which they are associated with. Its non-invasiveness and high temporal resolution along with its ease of setup and relatively low cost have established surface electroencephalography (EEG) as the most widely used brain imaging method for BCI (Lotte et al., 2018). While BCI was initially developed within the confines of standard laboratory conditions, recent advances in mobile neurophysiological sensing devices and artificial intelligence have led to a renewed interest in BCI applied to real-world contexts (Fairclough and Lotte, 2020). As a result, these neurotechnologies are now expanding in the clinical field (assistive technologies, motor rehabilitation, etc.), spreading to the entertainment industry to enhance gaming experience, but also extending to the general public through well-being (*e.g*. meditation and relaxation induction, sleep improvement) and domotics (*e.g*. home automation) applications (Brouwer, 2021). Following this trend, the range of neuroergonomics applications that can benefit from BCI broadens as sensors and interfaces become ever less intrusive (Gramann et al., 2021; Dehais et al., 2020a). Indeed, BCI has the potential to alleviate mental and physical loads associated with the repetition of straining actions (Carelli et al., 2017; Maksimenko et al., 2018), to improve task performance both in terms of its precision and speed, and to promote new forms of interactions to enhance human-machine teaming (Dehais et al., 2020b). Such assistance is particularly desirable in the context of aircraft operations. Flying a plane is a highly demanding task from a cognitive standpoint since it takes place in a dynamic and uncertain environment. It is well established in cognitive science literature that attentional resources are limited (Kahneman, 1973). This limited capacity implies that the pool of cognitive resources has to be distributed across competing sensory modalities (Wickens, 2008). The amount of cognitive resources available at any given time is also mediated by other factors such as stress and mental fatigue. Following prolonged periods requiring attentional focus, individuals typically exhibit performance decline as fatigue ensues (Dehais et al., 2018). Pilots have to attend to and assess several sources of visual and auditory information spread around the cockpit, and have to take decisions under time pressure and execute maneuvers in a timely manner (Behrend and Dehais, 2020; Wickens and Dehais, 2019). The combination of the stress-inducing context of flight operations and the need to sustain attentional focus over long periods of time is particularly taxing and tiring for pilots. The accumulation of mental fatigue and stress hinders access to the pool of cognitive resources (Hancock et al., 2006), which in turn can leads to completely overlooking information (*e.g*.., attentional tunneling). This phenomenon has been observed in experienced pilots and can have dramatic consequences (Wickens and Alexander, 2009; Dehais et al., 2019a; Mumaw et al., 2019). The on-line estimation of pilots’ monitoring ability combined with implementing new types of interactions through neuro-adaptive technologies may therefore have critical implications for flight safety.

There are several ways whereby BCI can enhance pilot-cockpit teaming. Firstly, active and reactive BCIs allow users to perform interactions under voluntary control via their brain waves (Lotte and Roy, 2019; Hong and Khan, 2017). Several studies disclosed that such technology can be used by pilot to control the flight-path of airplanes and drones (Nourmohammadi et al., 2018; Rodriguez-Bermudez et al., 2019; Fricke et al., 2014). Active BCI require the users to deliberately produce brain signals to interact with the BCI, as with mental imagery. In contrast, reactive BCI (rBCI) makes advantage of the user’s cerebral responses elicited by different stimuli. Each stimuli is associated with a different command. Due to their robustness and quick onset (*i.e*., below 50 ms), the Visual Evoked Potentials (VEP) elicited through the presentation of modulated visual stimuli are a popular and efficient approach for rBCI (Zhu et al., 2010; Chevallier et al., 2021). They offer very high classification performance (Nakanishi et al., 2018; Nagel and Spüler, 2019). Neural activity recorded in the visual cortex with surface EEG is sensitive to temporal and frequency features of the visual stimuli. Two main types of VEP-based paradigms can be distinguished: Steady-States Visually Evoked Potentials (SSVEP) and code-Visually Evoked Potentials (c-VEP). While SSVEP consists of the periodic modulation of visual features (*e.g*., contrast, color) at a regular frequency sustained over time, c-VEP waveforms are generated by pseudo-random binary (on/off) sequences. The c-VEP pseudo-random sequences are usually broadband and aperiodic (Shirzhiyan et al., 2019). They are defined so that temporal shifts have minimum cross-correlation and ensure a good separation between classes. VEP-based paradigms present the main advantage of relatively short training time, compared to P300 ERP or Motor Imagery paradigms, as only a low number of short-lasting trials is usually required to achieve accurate calibration (Nagel and Spü ler, 2019). In the field of aviation, visual BCI could offer promising perspectives for pilots by allowing them to free their hands when interacting with some actuators (*e.g*., landing gear, flaps). This could be particularly relevant during high g-force scenarios or critical flight phases (*e.g*. low altitude situations) that require both hands to control the stick and the thrust.

A second approach to improve pilot-cockpit teaming is to consider the use of passive BCIs (pBCI). This latter type of neuroadaptive technology supports implicit interaction by monitoring mental states (*e.g*., stress, fatigue) and adapting human-machine interactions to overcome cognitive bottlenecks (Zander and Kothe, 2011; Ewing et al., 2016). Several pBCI studies have been implemented in the field of aviation to infer mental workload (Dehais et al., 2019a; Gateau et al., 2018), failure of attention (Dehais et al., 2019b,c), flying performance (Scholl et al., 2016; Klaproth et al., 2020), and mental fatigue (Dehais et al., 2018). Interestingly enough, some authors managed to close the loop by triggering adaptive automation to prevent mental overload and task disengagement (Prinzel et al., 2000; Aricò et al., 2016). Generally, specific frequency-domain features are computed over the electrophysiological signal to account for different mental states (for a review, see Borghini et al. (2014)). For instance, changes in mental demand are related to the variation in the alpha band power and in theta band power over fronto-parietal sites (for a review, see Borghini et al. (2014)). Some studies also disclosed that increased beta (Matthews et al., 2017) is a neural marker of higher mental efforts. Alternatively, time-domain analyses over the EEG signal (*i.e*., event-related potential) can also predict variations of cognitive performance and attentional states (Roy et al., 2016; Brouwer et al., 2012; Dehais et al., 2018). One major drawback of such approach is that the calibration requires the induction of the mental states (*e.g*. different levels of stress or attention) in a repetitive fashion to train the model. It is difficult to achieve under laboratory conditions and, more importantly, is detrimental to the user experience. One alternative approach would be to take advantage of VEP and to use code-VEP tagging stimuli to implement a pBCI. By placing these flickers within the background of different regions of interest, one can measure the intensity of the brain response and derive the level of attention allocated to these specific areas.

Taken together, all these studies demonstrate the benefit of rBCI and pBCI to improve pilot-cockpit teaming and flight safety. However the rBCI and pBCI technologies to date have been used separately, whereas many of everyday-life tasks involved conjointly some voluntary interactions with a user interface and the monitoring of the state of the machine. Moreover, the same device (*e.g*. EEG) could be used to collect brain data and feed different algorithms in charge to control an interface and to infer the user mental state. Such an approach would pave the way to design a novel concept of neuroadaptive technology, namely a dual BCI (dBCI). By “dual”, we mean that it combines both “reactive” and “passive” components of the BCI to support direct and implicit interactions for end-users. This approach echoes with the concept of invasive bidirectional BCI for disable people that have been designed to translate motor cortex activity into signals to control an apparatus and to provide feedback by translating artificial sensor to restore sense of touch (Hughes et al., 2020). In our case, the dBCI relies on a non-invasive technology such as surface electrodes placed over the scalp *(i)* to allow an end-user to directly control a machine by their brainwave and *(ii)* to allow a machine to communicate feedback to its end-user and to adapt human-machine interaction to overcome cognitive bottleneck.

The objective of this study was to implement such a dBCI in the cockpit. To meet this goal, we used the NextMind 9-electrode dry EEG system and their Unity framework (www.next-mind.com) that allows to implement asynchronous code-VEP based BCI for up to 10-class problems. We chose the NextMind as their hardware is light, fast to setup and the classification is plug-and-play but this could have been done with another BCI system. Our main motivation was not to focus on the algorithmic implementation but to demonstrate the effectiveness of our genuine dBCI to decode pilots’ intention to interact with the flightdeck (reactive, rBCI) and to infer their level of attention on a monitoring task and to adapt the interaction accordingly (passive, pBCI) - (see Figure 1.

**Figure 1.**
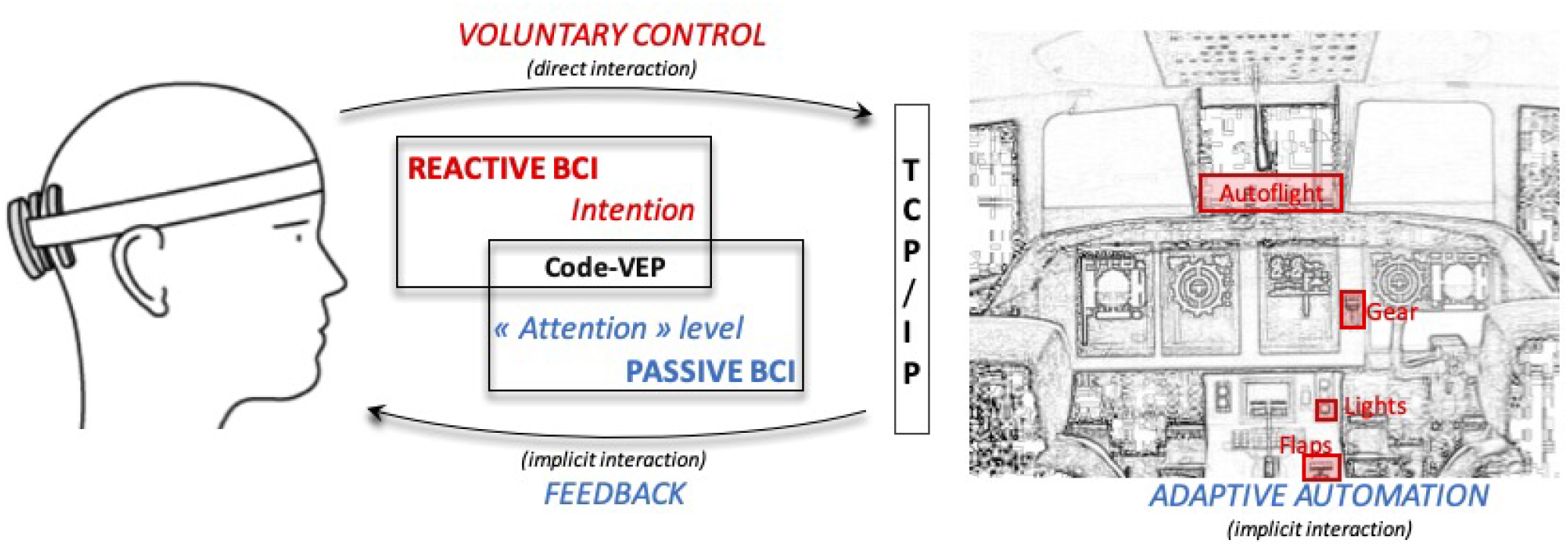
Implementation of our dual passive and reactive Brain Computer Interface in the cockpit. It enables human-machine bi-directional communication: **(1)** the pilots can directly interact with some flight deck actuators (*e.g*.., the landing gears - as shown by the red rectangles) using information from the EEG signal, and **(2)** the flight deck can send visual feedback to the pilot and adapt the interaction when poor cognitive performance is detected.

The designed scenario in the flight simulator requires to tackle both direct (rBCI) and implicit (pBCI) interactions. Indeed, the participants had to perform several checklists operated through rBCI and a radar monitoring task which was handled by the pBCI while completing an engaging traffic pattern exercise. We evaluated the rBCI during two contrasted experimental conditions in terms of task difficulty. In one condition, the participants were interacting with the rBCI alone while the plane was operated by the autopilot. In another condition, they were operating the rBCI while flying the plane and monitoring the radar. We collected objective measures (reaction time and accuracy) and subjective measures (level of mental demand and frustration). It was expected that the multitasking condition would lead to longer reaction times to interact with the rBCI, more frustration and lower accuracy. We evaluated the pBCI component using classical machine learning metrics (*F*_1_ score, sensitivity and specificity).

## 2 MATERIAL AND METHOD

### 2.1 Participants

Twelve participants (mean age = 29.8 year old, mean flight hours = 488.9), all students and staff members from our aeronautical university, took part in the experiment. The study was approved by the local ethics committee (approval number 2020 − 334). We followed specific COVID procedures implemented by our Health, Safety and Working Conditions Committee (anti-COVID face masks, sanitization of the flight simulator, disinfection of the eye tracker and EEG device following a thorough cleaning procedure).

### 2.2 Dual BCI implementation

We used the NextMind Unity Software Development Toolkit (SDK) ^1^ to implement the rBCI and the pBCI. An MSI laptop (processor Intel *i*7 − 6700*HQ* with 16*Gb* RAM and NVIDIA GeForce GTX 960*M* graphic card) was used to received the EEG data stream trough Bluetooth Low Energy. This PC executed the decoding algorithms and sent the decoded brain command to the core simulator via socket TCP/IP (using an Ethernet connection) that consisted of a thread of four commands (integer type) to change the states of the flaps, the landing gear, the landing and taxi lights and the autopilot (see Figure 1 and Figure 2).

**Figure 2.**
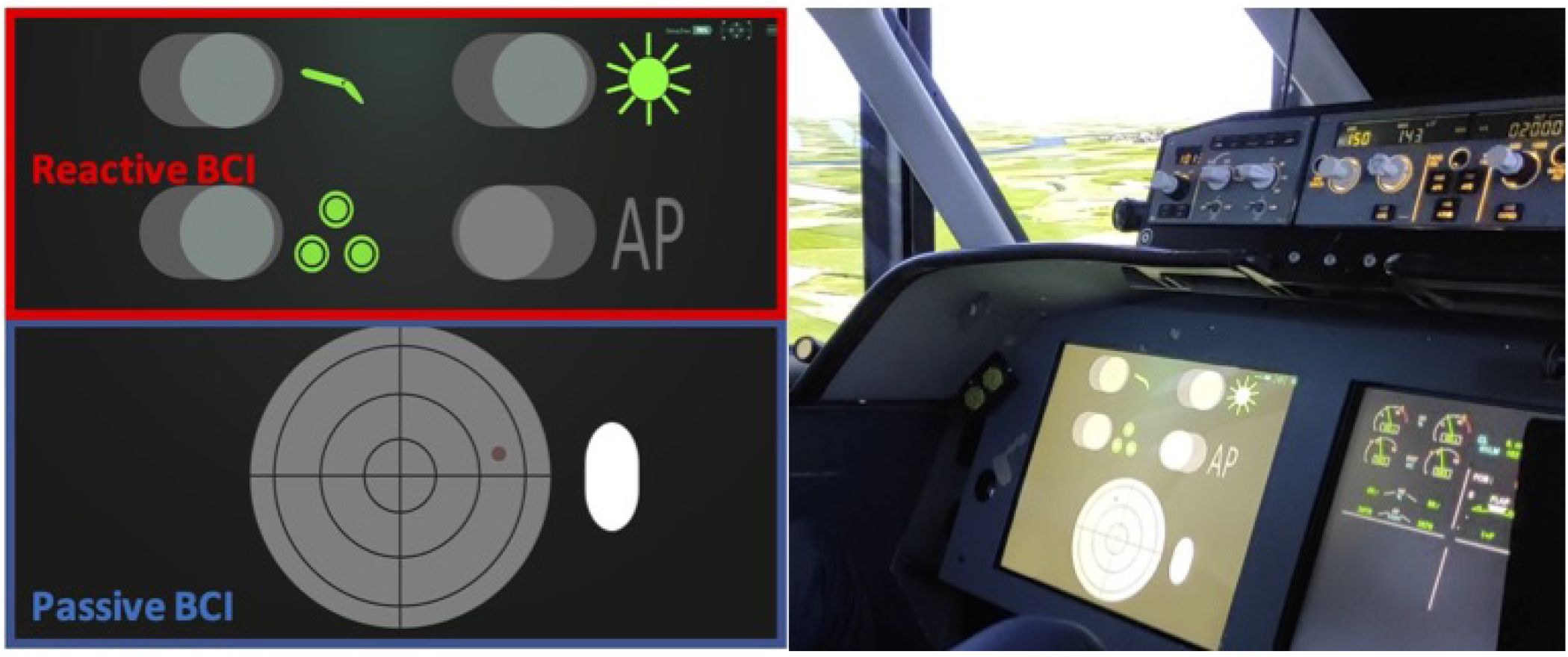
**Left**: the participants had to perform the checklists with the reactive BCI (in red, upper part of the screen) while their monitoring performance on the radar screen was assessed with the passive BCI (in blue, lower part of the screen). **Right**: the dual BCI setup in the flight simulator.

**Figure 3.**
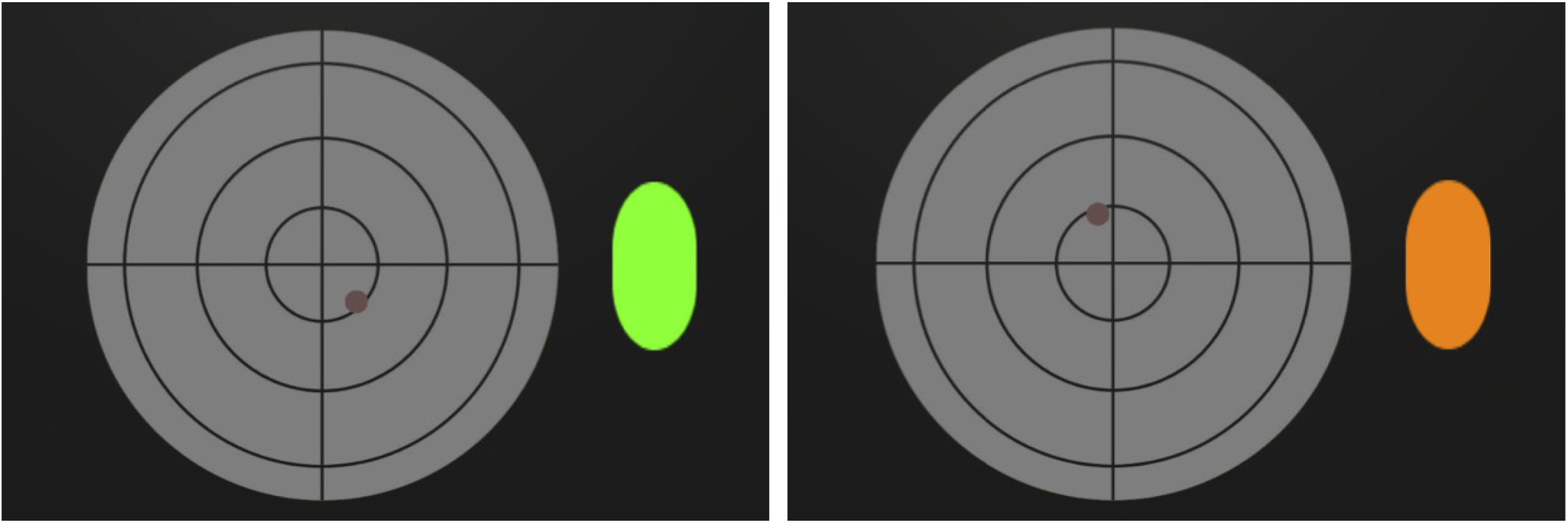
Left: the pBCI detected that the pilot was correctly monitoring the radar and, in return, displayed a green visual feedback. Right: the pBCI detected that the pilot did not notice the potential collision. It then displayed an orange warning and automatically simulated the avoidance maneuver.

#### 2.2.1 rBCI

The rBCI display consisted of four toggle switches (diameter = 4*cm*) that are defined as “NeuroTag” interactive elements (*i.e*., a NextMind code-VEP) in Unity. They are depicted in the upper left part of Figure 1, outlined in red. These switches are dedicated to lower/retract the flaps, switch on/switch off the taxi and landing lights, to lower/retract the landing gears and to engage/disengage the autopilot system (see Figure 2 - left).

#### 2.2.2 pBCI

For the pBCI, the region of interest was a radar. We have integrated a “NeuroTag” (diameter = 13.1*cm*) in the background. It is depicted in the lower left part of Figure 1, outlined in blue. The brain response to this background c-VEP was used to assess the level of attention of the pilot to the region. Thus, instead of using the regular rBCI classification procedure, we have taken advantage of a continuous *confidence score* (∈ [0, 1]) defined by the NextMind Engine as a proxy measure for attention to the radar. This score reflects the classification certainty for a rBCI. If this score was too low and a collision was incoming, our system would infer that the pilot is not paying attention at the radar and therefore not able to avoid the collision. It would then automatically activate the anti-collision maneuver and trigger an orange visual alert. Based on a preliminary experiment with 5 participants, we set a threshold of 0.1 on the *confidence score* to have good responsiveness to determine whether the pilots were actually monitoring the anti-collision radar or not (see Figure 2 - right). The classification processes are different between the rBCI (regular and hard decision of classification) and the pBCI (the certainty of classification by the model as attention probe), though they both use the response to the same type of c-VEP as input.

### 2.3 Flight simulator

We used our three-axis hydraulic (pitch, roll, height) motion flight simulator to conduct the experiments. It has eight external panoramic displays that reproduces the outside world based on the Flight-Gear open-source software^2^. It simulates twin-engine aircraft equipped with two side-sticks, a thrust, two rudders, and an advanced auto flight system (Figure 2). Its user interface is composed of a Primary Flight Display, a Navigation Display, and an Electronic Central Aircraft Monitoring Display and a head-up-display (HUD). The HUD provides basic flight parameters (speed vector, angle of attack, total energy) and thus allows the pilots to control the flight path and the speed of the airplane. The flight simulator refreshes every 20*ms* the inputs from the BCI system send trough the TCP/IP connection, in addition to the information coming from the traditional organs of piloting like the autopilot button, the flaps and landing gear levers, *etc*. The dual-BCI user interface (checklists and anti-collision radar) was displayed on a head down 17^′′^ screen (1280 × 1024 - 60 Hz) facing the participants (head distance = 80 cm).

### 2.4 Scenario

We have designed a scenario in which the participants had to perform two checklists (rBCI) and a radar monitoring task (pBCI). The flying task consists of a first landing at Toulouse Blagnac runway (33R) followed by a go-around, then a traffic pattern exercise leading to a final landing. The participants had to use the rBCI during the:

1. `first landing`: to lower the gear, to lower the flaps, to switch on the landing and taxi lights ;
2. `go-around`: to retract the gear, to retract the flaps, to switch off the taxi and landing lights and to engage the autopilot;
3. `crosswind`: to disconnect the autopilot;
4. `downwind`: to lower the flaps and to switch on the lights;
5. `final landing`: to lower the landing gear.

Meanwhile and while flying the plane, participants should monitor the anti-collision radar (pBCI). The anti-collision task was not performed in the auto-pilot condition (low workload) but only while flying (high workload). The collisions were not linked to the actual flight pattern exercise and only appeared on the radar. In practice, it implies looking frequently at this radar to determine if another plane (represented by a red circle) reaches the center of the radar, indicating a collision. In the case of an incoming collision, pilots were asked to focus on the radar. This would mean that they have acknowledged it and would avoid it. When a collision is incoming, an anti-collision system would be triggered by the pBCI either by *i)* the detection of sufficient attention to the radar *ii)* or by the detection of inattention of the pilot. With this scenario, the expertise of the pilot to avoid collisions is primary and it is bypassed only if the system estimated that the pilot is distracted. It is different to a scenario that would systematically activate the anti-collision manoeuvre, which sets aside the pilot assessment of the situation. Thus, the idea of this pBCI is to offer supplementary assistance to the pilot during overwhelming situations but not to automate his or her tasks.

In order to assess the effect of mental workload and multitasking on the use of rBCI, we manipulated two variations of the experimental conditions:

- `rBCI alone condition (single task condition)`: the participants only had to perform the different checklists without flying (the aircraft was in automated flight mode). They also did not have to do the anti-collision monitoring task;
- `r/pBCI and Flying condition (multi-tasks condition)`: the participants had to perform the checklists and the radar monitoring task while manually flying to perform the five legs of the scenario.

These stand for two realistic conditions of flight: auto-pilot mode and full control.

### 2.5 Protocol

The participants underwent 30 minutes of training for the flight simulator without the dBCI. It included a short tutorial about how the simulator worked (user interface, flight parameters), several landings and a complete traffic pattern exercise. Participants were then equipped with the 9-electrode NextMind EEG headset (Oz, PO7, PO8, PO1, PO3, PO4, P3, P4), placed over the visual cortex with the lowest electrode (Oz) set over the inion, as recommended by the manufacturer best practices ^3^. Following this, participants went through the BCI calibration phase which output a calibration score (between 1 and 5). During the calibration phase, the participant had to focus on a single c-VEP for 40 seconds. The calibration was redone if the score was below 4 to ensure satisfactory performances. After this setup, participants completed the two conditions (rBCI alone and r/pBCI + flying). The order of the two conditions was pseudo-randomized to counterbalance between all participants and to control for potential fatigue effects: six participants started with the “rBCI alone” condition and the remaining six with “r/pBCI and flying” condition. Finally, after completing each experimental condition, participants were asked to fill the subjective questionnaire. The total length of the experiment for a subject was about one hour.

### 2.6 Measurements

#### 2.6.1 Subjective measures

The participants reported their subjective levels of mental effort and frustration to perform the checklists using a Likert scale (1 = very low, 10 = very high) in the two experimental conditions. These subjective assessments were done immediately after completing the experiments.

#### 2.6.2 Objective measures

We used *a posteriori* the recording from a Tobii Glasses II eye tracking system (Tobii Pro AB, Stockholm, Sweden) to manually evaluate the efficiency of the dBCI system. Tobii Glasses II is a wearable eye tracker with an embedded scene camera. The recording consists in the video of the scene camera, with a large angle, with in overlay the gaze point of the subject, estimated by the Tobii algorithm. The eye tracker is not part of the dBCI system. The recording was used *i)* to compute the accuracy of the rBCI and measure the reaction time to perform the checklist events, and *ii)* to quantify the performance of the pBCI. We have manually checked and timed when the user’s gaze entered a check-list item or the radar and the activation time, or false positive. For the pBCI we considered that a collision was missed if in the recording the gaze of the subject was not on the radar when the point reached the center.

## 3 RESULTS

In this section, we present the subjective and quantitative results of our dual (reactive and passive) BCI.

### 3.1 Subjective results

A paired t-test (*p <* 0.001) demonstrated that the mean level of frustration to interact with the checklist items was significantly lower in the “rBCI alone” condition (mean = 2.6, SD = 2.0) than in the “r/pBCI and flying” condition (mean = 4, SD = 2.2). Similarly, a paired t-test (*p <* 0.001) demonstrated that the mean level of mental workload to interact with the checklist items was significantly lower in the “rBCI alone” condition (mean = 2.2, SD = 1.4) than in the “rBCI and flying” condition (mean = 5, SD = 2.0) - see Figure 4. Such reported results were expected as the “rBCI and flying” is way more demanding compared to ‘rBCI alone”. We selected these two conditions as they represent two classical flight situations: auto-pilot with small workload and flying with high workload.

**Figure 4.**
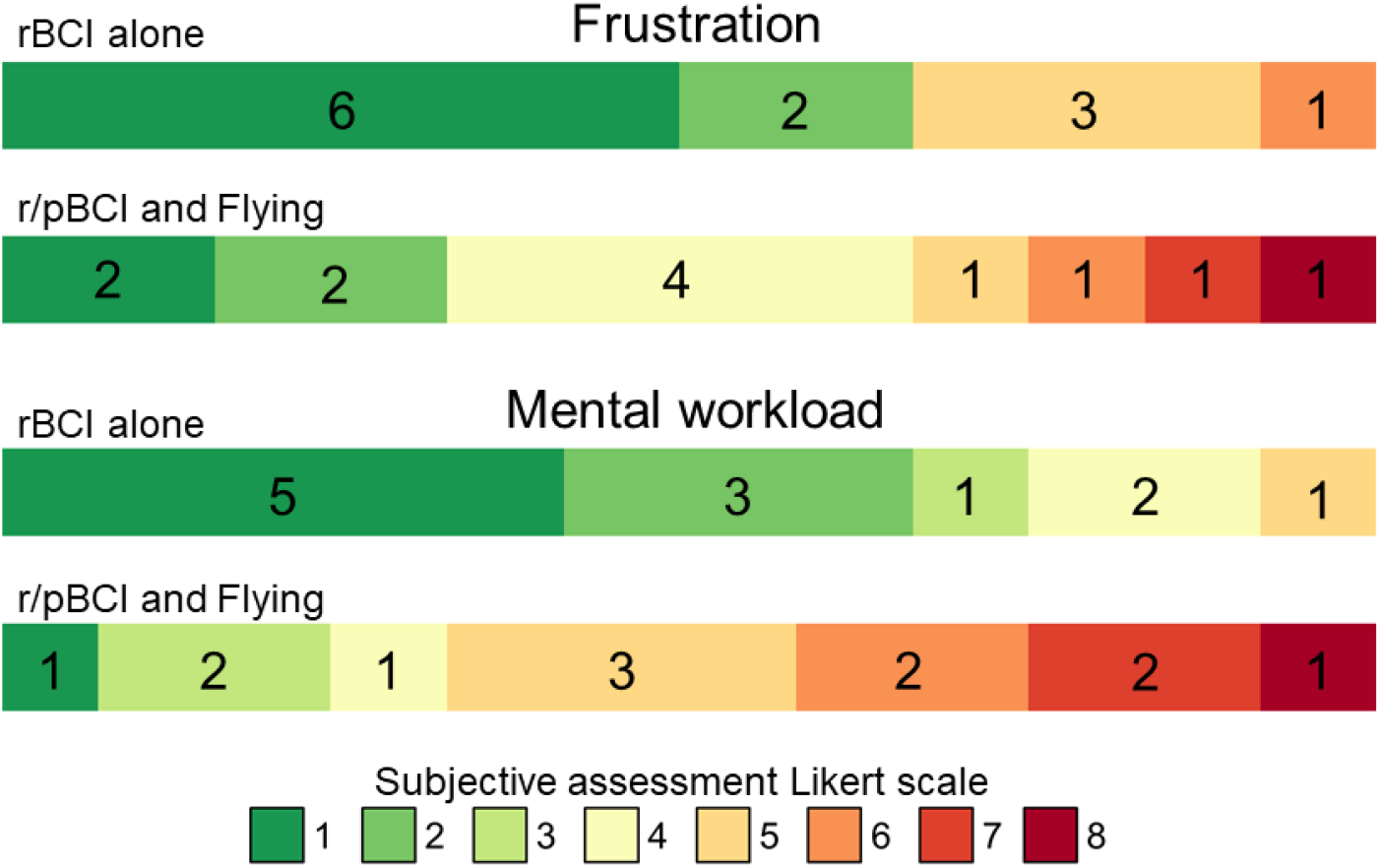
Subjective results using a Likert scale for the level of frustration and mental workload between the two experimental conditions.

### 3.2 rBCI objective results

In the “rBCI alone” condition, the classification accuracy reached 100% since all the participants interacted successfully with the checklist items without experiencing any false positives (*i.e*., activation of an undesired item). Unlike a traditional rBCI with a fixed epoch length, here the epoch length varies. In the “r/pBCI and flying” condition, all the participants managed to fly the different legs of the aircraft while interacting with the different checklist items. The classification accuracy reached 98.5% since only two single false positives occurred out of 132 trials (*i.e*., 11 checklist items × 12 pilots).

A paired t-test indicated that the mean reaction time to activate the checklist items was significantly lower (*p* = 0.002) when interacting with the rBCI alone (mean = 1643.7*ms*, SD = 390.6 ms) than when interacting with the rBCI while flying and monitoring the radar task (mean = 2493.1*ms*, SD = 1055.4*ms*). Mean reaction times per participant can be found in Figure 5. The variability in reaction times between users is quite limited when the rBCI is operated alone and is larger when operated as a secondary task to flying and radar monitoring (pBCI). It is well known that the performance of a BCI are tightly related to the subjective mental workload experienced by the users (Felton et al., 2012), the higher the less the BCI performance would be. While the task difficulty is the same for all subjects, the subjective workloads experienced are different as it depends also on the skills, fatigue, *etc*. of the subjects. To better quantify the variability, some further studies with more items per pilot and more pilots could be conducted.

**Figure 5.**
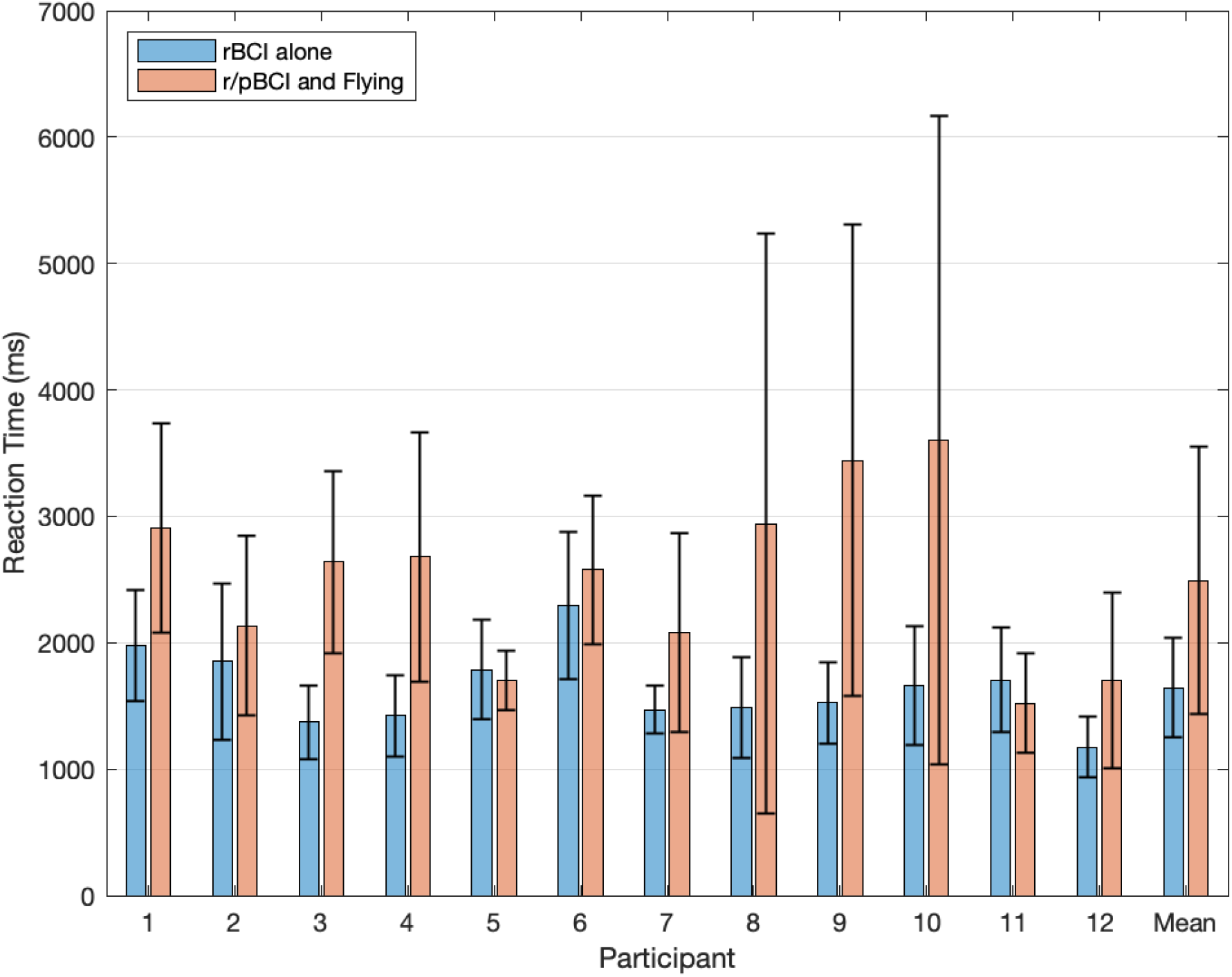
Mean reaction time per participant and grand average reaction time to activate the checklist items when interacting with the rBCI alone (in blue) and when interacting with the rBCI along with the flying task and the pBCI (in orange)

### 3.3 pBCI objective results

The pBCI system have classified as missed by the pilots, a total of 53 potential collisions out of 203 events. Among these 53 missed collisions, the recordings from the eye tracker, manually examined, showed us that in 9 cases, the pilot was actually fixating the radar but the pBCI considered as an attentional lapses (False Negative, FN). For the remaining 44 cases (True Negative, TN) and still based on the manual study of the recordings from the eye tracker, the pBCI had accurately detected that the pilot was distracted and compensated attention errors by automatically triggering the orange alert. During the *post hoc* analysis, we have also determined that in 8 cases among the 150 potential collisions classified by the pBCI as acknowledged by the pilots, but the eye tracker disclosed that the participants were not actually paying attention (False Positive, FP). Therefore it makes a total of 150 − 8 = 142 true positives (TP). As such, the True Positive Rate (TPR, also called recall=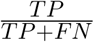) of the pBCI system was 94.04%. The precision 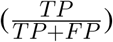 was 94.67% and it gives a *F*_1_-score (harmonic mean of precision and recall) of 0.94. The *F*_1_-score reflects the number of false positives and negatives along the true positives and negatives while traditional classification accuracy only provides information about true positives and negatives. Within our framework, missing a collision could have dramatic consequences, thus it was more informative to consider *F*_1_-score than accuracy. The True Negative Rate (TNR, also called selectivity=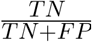) was 84.61%.

## 4 DISCUSSION

This study is the first demonstration of a dBCI system that promotes direct (rBCI) and implicit (pBCI) interaction between the user and the interface. In our task, the reactive and passive aspect did not interact together but were motivated by the task involving interactions with the flight deck and the monitoring of the radar screen. More specifically, the rBCI component allowed the participants to interact with the flight simulator, sending commands trough a TCP/IP connection, to change the state of specific flight deck actuators. This approach differs from previous aviation-oriented BCI studies, which apply BCI to directly control the aircraft’s trajectory (Nourmohammadi et al., 2018; Rodriguez-Bermudez et al., 2019; Fricke et al., 2014). It is important to note that in all these previous studies, the pilots were required to fully allocate attentional resources towards the BCI interface in order to perform the flying task whereas the flight performance under the BCI condition does not meet the standards of manual flying accuracy. Another important consideration is that operating an aircraft requires to constantly monitor a rich flow of information distributed across the cockpit. Therefore such artificial setups in which pilots only pay attention to a single source of information does not accurately reflect real-flight situations and may not be transferable to these use cases. Moreover, any relevant secondary task (*e.g*., radar monitoring, communication with air traffic control) or critical stimuli (*e.g*. alarms) will distract them from flying. The pBCI component was precisely dedicated to assist the pilot when performing a secondary task, namely monitoring an anti-collision radar task. To the authors’ best knowledge, this is the first time that a VEP is used to monitor attention. Traditionally, this is performed using frequency and/or time-domain features to assess the level of attention (Dehais et al., 2018; Brouwer et al., 2012).

The rBCI analyses disclosed state of the art results in the control condition (auto-pilot) with an accuracy of 100% and a average reaction time of 1.6*s*. It should be compared with other asynchronous, online and with 4 classes BCIs of the literature. Kalunga *et al*. (Kalunga et al., 2016) have reported a reaction time of 1.1 for a 3 class problem and an accuracy of 87.3% for they asynchronous and online BCI. In Gembler et al. (2020), authors have achieved a reaction time of 2.31*s* for a 4 class problem and a mean accuracy of 94.4%, also online and asychronous. The rBCI also led to very high accuracy (98.5%) in the condition where the participants had to operate it while flying the plane and monitoring the radar. The mean reaction was longer but quite acceptable (2.5*s*) and allowed the pilots to perform the checklist in a timely manner while still being able to navigate and safely land the plane. This slight decrease in classification accuracy (−1.5%) and increase in reaction time (+0.9*s*) can be explained in terms of higher mental demand. Indeed, the participants were engaged in a complex multitasking activity, leading them to divide their visual attentional resources between the flying, the radar, and the rBCI tasks. Our subjective results seemed to confirm this hypothesis as our participants expressed significantly higher mental workload and higher frustration in the multitasking condition. These results are consistent with a previous rBCI study reporting a 20% lower accuracy classification in the multitasking condition compared to the control condition (Vecchiato et al., 2016). Nevertheless, the implemented rBCI demonstrate sufficient efficiency and responsiveness to be operated, even while flying and with high workload which was the aim.

Beside that, the pBCI findings also seemed to indicate the soundness of VEP to probe the level of attention to a monitoring task. One of the main advantage of this approach is that the same calibration, lasting only 40*s*, was used to train the rBCI and the pBCI. To the best of our knowledge, this is the first online demonstration that flickering stimuli could be used for pBCI purpose. The behavioral results showed that our pilots missed a total of 44 out of 203 collisions. The pBCI provided assistance to the pilots by simulating safety maneuvers with an acceptable rate of false negatives. We believe that this hybrid approach provides flexibility since the expertise of the pilot is kept at the center of the design, while providing a safety net in case the pilot’s attentional resources are engaged on other aircraft operations. We consider it as an improvement compared to automatically activating the anti-collision safety whenever a collision is coming. In some cases, the pilot could have paid attention to this incoming danger but determined that the anti-collision safety was not necessary. In our scenario, the anti-collision safety was triggered only if the pilot deliberately chose to do it or if he or she was not paying attention. However, it is worthy to note that such adaptive automation could cause some drawbacks such as over-reliance on fallback mechanisms.

Despite these promising results it appeared that in eight cases out of 150, the pBCI did not detect pilot’s attentional lapses and failed to trigger the automatic maneuver (false positive). The occurrence of such failure could lead to critical scenarios in real-life situations. One has to keep in mind that we used a an empirical threshold set to 0.1 for all participants to infer the attentional state and trigger the pBCI. This fixed threshold, while proving to be efficient overall may not be optimal for each individual. A possible solution is to define an individualized threshold to obtain an optimal ratio between false positive and true negative. Interestingly enough, the findings disclosed that in nine instances out of 150 events, the pBCI classified that the participants detected the collision whereas the participants were not glancing at the radar as attested by the eye tracker (false negative). However, during the debriefing, the participants declared that they actually spotted the collision via peripheral vision. Such situation is possible since peripheral vision, via the rods, is highly sensitive to motion and flickering objects such as the VEP stimuli that were used in this experiment. For instance, some authors demonstrated the potential of using concurrent presentation of VEPs-based stimuli to assess processes outside of the focus of spatial attention (see for reviews Vialatte et al. (2010); Norcia et al. (2015)). Future work should investigate this hypothesis.

Following on these promising results, this work calls for refinement through an actual implementation of the dBCI in the wild, *i.e*. under operational conditions. For instance and from an operating time point of view, it is blatant that using one’s hands is still faster than the rBCI to interact with the cockpit. One solution is to provide hybrid interaction so that the pilots can choose to use either the BCI or physical modalities depending on the flight phase and time pressure. One could imagine that the pilot would interact trough the BCI when the autopilot is engaged, so to increase his comfort and availability, and manually during more dynamic phases of the flights (*i.e*., maneuvers), when full commitment is required. In the present study the eye-tracking data were used as to provide a ground truth to assess both passive and reactive BCI performances in *post hoc* analyses. The gaze data could also have been leveraged as a complementary source of spatial and temporal information to improve BCI speed and accuracy. Indeed, the eye-tracking data could provide contextual information, as to which area of the environment the user focus attention. This contextual information (regions of interest in eye-tracking terminology), would lead to the activation of only a subset of the VEP stimuli based on the localization of user’s attention (Lin et al., 2019). This approach would allow to artificially increase the number of classes while using a constant number of distinct VEPs therefore reducing the complexity of the classification problem (Stawicki et al., 2017). This lower number of distinct VEP also implies to acquire less calibration data compared to traditional paradigms in which each class is represented by a distinct VEP. Moreover, by triggering stimuli presentation, the eye-tracking data would provide the precise onset time of the stimuli thus giving valuable information for asynchronous BCI allowing to compute classification only when necessary and optimize computing power and time requirements. Additionally, gaze contingent rBCI would reduce the bottom-up influences of the visual stimuli on the user attentional resources. Indeed, the high contrast nature of stimuli commonly used in VEP-based paradigms may distract attentional resources away from the primary task (Zhao et al., 2018). From an user experience perspective, the reduction in the number of VEP stimuli presented at the same time may improve visual comfort. Eventually, visual fixations on the radar area may be used as a two-step certification process to validate the VEP-based pBCI decision. Similarly for the rBCI, visual fixations within VEP stimuli area could be used to confirm the intention of an user to interact with a command. Overall eye-tracking information concurrent to VEP-based BCI classification outputs may be used to further improve classification performance and reduce the rate of false positives. This latter point is particularly important in the context of translating the proposed dual BCI system to real cockpit day-to-day flight operations as meeting high standards of aviation certification criteria is particularly challenging (10^−3^ allowable failure probability). In the context of aviation, any undesired activation of a command could jeopardize flight safety.

To conclude, we believe that the concept of dBCI opens promising prospects to improve human machine symbiosis for neuroergonomics applications in many domains such as transportation, industrial workplaces, medical care but also for disable people. We have developed a proof of concept that relies on the code-VEP based stimuli and classification tools provided by the NextMind company. However, this approach is limited as their system is a black-box. The EEG data streams are not accessible and the algorithms are not provided by the manufacturer. Thus, we hope that this study should encourage the development of open-source c-VEP code for the scientific community. Alternatively, other types of stimuli and classification procedures could be explored. For instance, the decoding of user’s expectation (Zander et al., 2016) represent an interesting alternative since it does not require to present additional stimuli in the user interface. Combining together active and reactive tasks and the use of hybrid EEG and functional near infrared spectroscopy based BCI could maximise the number of commands (Hong and Khan, 2017) provided by a BCI. Regarding the pBCI, the use of more traditional features (*e.g*. changes in the well defined band of power for EEG) or more advanced ones like brain connectivity could be considered to target specific degraded mental states (cognitive fatigue, failure of auditory attention) so as to trigger the most appropriate neuro-adaptive solutions (Dehais et al., 2020b). Indeed, several solutions could be designed to dynamically optimize human-machine teaming by *(1)* adapting the user interface using notifications to alert of impeding hazards, *(2)* adapting the task and the level of automation to maintain the performance efficiency of the operators, and *(3)* “neuro-adaptating” the end-users to warn them of their mental state and stimulate them to augment performance (*e.g*. neurofeedback). We truthfully hope that this study will foster research efforts to improve the concept of dual BCI for safer, seamless and efficient human-human and human(s)-machine(s) interactions.

## STATEMENTS

### Author Contributions

FD contributed to the conception and the design of experimental paradigm. FD, SL, NT-V recorded the data. FD, NT-V, SL, LD performed part of the analyses. SV, PL, GF, JTT implemented the experimental set-up in the flight simulator. FD, SL, LD wrote the manuscript and all authors revised the manuscript. All authors read and approved the submitted version.

## Acknowledgments

The authors wish to sincerely thank all the pilots who participated in the experiment. The authors would also like to acknowledge the Artificial and Natural Intelligence Toulouse Institute (ANITI) and the Axa Research fund Neuroergonomics chair for flight safety for currently funding FD.

## Footnotes

1 https://www.next-mind.com/developer

2 www.flightgear.org

3 https://www.next-mind.com/documentation/sensor-manual/

